# Chart Builder: An Interactive Tool for User Driven Data Visualization in the Electron Microscopy Data Bank

**DOI:** 10.64898/2025.12.03.691848

**Authors:** Neli Fonseca, Amudha Kumari Duraisamy, Zhe Wang, Sriram Somasundharam, Minoosadat Tayebinia, Lucas C. de Oliveira, Miao Ma, Jack Turner, Ardan Patwardhan, Gerard J. Kleywegt, Matthew Hartley, Kyle L. Morris

## Abstract

The cryogenic sample-electron microscopy (cryoEM) field has generated significant amounts of 3D Electron Microscopy (3DEM) volumetric data and associated metadata, now comprehensively archived in the Electron Microscopy Data Bank (EMDB - www.emdatabank.org) and the Electron Microscopy Public Image Archive (EMPIAR - www.empiar.org). Harnessing the full potential of these resources requires robust, flexible, and publicly accessible tools for data exploration, analysis and retrieval. Here, we present Chart Builder, an interactive web-based platform that enables researchers to create customizable, publication-quality visualizations directly from archival metadata, validation assessments, and cross-reference annotations. Chart Builder integrates the same query-driven and flexible Solr search system as EMDB search, into a user interface with tools to assist users to filter, group, and compare data without programming expertise. It supports multiple chart types (including line, bar, area, scatter (2D and 3D), histogram, bubble, pie, geographic and Venn diagrams) with customizable axes, data series, and statistical operators. Users can apply global filters, define temporal, categorical, or custom query-based axes, and explore multi-dimensional relationships interactively. Chart data-points are linked to their underlying datasets, such that visualisation interaction opens entry-level or archive-level search results for inspection and datasets from charts may be exported in several ways. Findability, accessibility, interoperability and reusability of data are facilitated by these direct access and export mechanisms, including HTML embedding, persistent URL sharing and chart/data download options. By combining interactivity and ease of use with up-to-date access to the EMDB and EMPIAR archive metadata, both computational and experimental communities may explore and visualize current metadata and export to formats for further analysis or as publication-ready figures. Chart Builder promotes community-driven data analysis and empowers users to evaluate trends in the biological 3DEM field. Chart Builder is freely accessible and fully integrated into the EMDB website at https://www.ebi.ac.uk/emdb/statistics/builder/.

## 1 Introduction

The Electron Microscopy Data Bank (EMDB) is a public repository for three-dimensional electron microscopy (3DEM) volumetric reconstructions from cryogenic sample-electron microscopy (cryoEM) and cryogenic sample-electron tomography (cryoET) (wwPDB Consortium, 2024). The Electron Microscopy Public Image Archive (EMPIAR) holds cryoEM and cryoET 2D and 3D image experimental and derived data, volume electron microscopy (vEM) and soft and hard X-ray tomography (XT) data (Iudin *et al*., 2023). Both are entry-based deposition databases, with over one hundred schema-controlled data fields describing individual entries. Together, EMDB and EMPIAR form a comprehensive resource ecosystem that represent data and metadata from experimental data collection, that data’s reduction and interpretation, through to the final 3DEM reconstruction of a biological target, predominantly via averaging techniques. Where a molecular reconstruction is targeted, these data may have model coordinates derived, which may be stored in the PDB database (wwPDB Consortium, 2019). The availability of data from these databases and connections between them creates scientific transparency, reproducibility, and innovation in cryoEM (Patwardhan *et al*., 2017) Importantly and more broadly this translates the same benefits to developments in experimental approaches (Kimanius et al., 2024, Jamali et al., 2024), structural knowledge (PDBe-KB consortium, 2020) and predictive structural biology data (Jumper *et al*., 2021, Varadi *et al*., 2022) made possible by the reuse of FAIR data.

The cryoEM field has been experiencing rapid growth, driven by rising investments in structural biology research, technological advances in throughput and quality of cryoEM imaging (Kühlbrandt *et al*., 2014) and the success of image processing softwares such as RELION (cite) and cryoSPARC (cite). Most often molecular structures are the targets of these studies leading to these data being archived in the core archives, EMDB, EMPIAR and PDB. This surge in data availability is highly valuable but presents mounting challenges for the community in terms of data analysis, quality assessment and underscores the essential need for accessible, standardised statistical summaries and visualization tools (Lawson *et al*., 2021).

Statistics plots have been available in the EMDB since its inception through the statistics page (formerly named EMstats), a collection of predefined charts developed in collaboration with the EM community. Initially containing a few dozen, it now includes over one hundred charts, summarizing the archives’ overall contents and trends (Tagari *et al*., 2002). These charts provide high-level insights into features such as resolution distribution, database growth over time, and methodological parameters (https://www.ebi.ac.uk/emdb/statistics). Additionally, for any set of search results, a set of nine dynamic plots can be viewed from the search statistics. The chart types are fixed but summarize the subset of retrieved entries. However, users wishing to generate any deeper statistical summaries or even analytical queries of the EMDB and EMPIAR are required to write bespoke scripts to access and retrieve the database content.

The motivation for this work is to provide an accessible, flexible, user-friendly, query-based visualization plotting tool designed to facilitate data exploration and enable the rapid creation of statistical and analytical enquiries without requiring coding skills or deep technical knowledge of the database. The EMDB Chart Builder extends and integrates EMDB REST API endpoints into an interactive, query-based framework that enables users to design customized visual summaries of EMDB data. It represents the next stage in the evolution of EMDB statistics, offering greater flexibility, transparency, and direct access to analytical tooling for dataset discovery and data-driven insights across a rapidly growing cryoEM archive.

## 2 Materials and Methods

### 2.1. Data sources and availability

The statistical visualizations in this work are based on four main data sources: author metadata provided during database submission to deposition systems either to EMDB via OneDep (Young et al., 2017) or to EMPIAR directly, and added-value annotations from EMDB EMICSS (Duraisamy *et al*., 2025) and EMDB Validation Analysis (Wang et al., 2022). Data from these sources are indexed in the EMDB Solr search engine and provided to the chart builder via access to public REST API endpoints. The user of the chart builder may identify metadata that they wish to explore and plot by searching all fields available in the search system (https://www.ebi.ac.uk/emdb/documentation/search/fields).

#### 2.1.1 EMDB metadata

The EMDB metadata is provided by the authors (depositors) during the deposition of cryoEM maps data and its associated metadata in the OneDep system (Young *et al*., 2017) through the OneDep deposition interface (https://deposit-pdbe.wwpdb.org/). Captured data includes but is not limited to key information describing each three-dimensional EM (3DEM) reconstruction, such as authors, dates, sample information, experimental information, map details and metadata for deposited files (Westbrook *et al*., 2022). These metadata are stored by the OneDep system in the mmCIF dictionary format (https://mmcif.wwpdb.org/) and translated into the EMDB schema controlled XML data model for dissemination (https://www.ebi.ac.uk/emdb/documentation#data_model).

#### 2.1.2 EMPIAR metadata

EMPIAR metadata is supplied by EMPIAR depositors, from the EMPIAR deposition system (https://www.ebi.ac.uk/empiar/deposition), describing administrative and technical data for raw imaging datasets, including contributor information, dataset descriptions, sample identifiers and cross references to EMDB.

#### 2.1.3 Validation Analysis metadata

The validation of EMDB entries provides standardized assessments of map and model quality using a broad set of metrics calculated by the Validation Analysis package (Wang et al., 2022). Key metrics that are made publicly available include resolution values from internally calculated (Chen *et al*., 2013) and deposited Fourier Shell Correlation (FSC) plots with multiple cutoffs, Q-score (Pintilie *et al*., 2020), Atom inclusion (Lagerstedt *et al*., 2013), CCC (Liebschner *et al*., 2019) and SMOC score (Joseph *et al*., 2017) for model-map fitting quality.

#### 2.1.4 Cross-reference annotations

Database cross-references delivering added-value annotations are systematically gathered by the EMICSS framework (Duraisamy *et al*., 2025). It generates comprehensive cross-references for each EMDB entry, providing direct links to a vast array of biological and structural databases. Specifically, it connects to resources related to **protein sequences**: UniProt (The UniProt Consortium, 2025), **ontological labels**: Gene Ontology (Aleksander *et al*., 2023); **structures**: PDB (wwPDB Consortium, 2019), PDBe-KB (PDBe-KB Consortium, 2022), AlphaFold DB (Varadi *et al*., 2022), EMPIAR (Iudin *et al*., 2023); **complexes**: Complex Portal (Meldal *et al*., 2015), **domains**: Pfam (Mistry *et al*., 2021), Rfam (Kalvari *et al*., 2021), InterPro (Paysan-Lafosse *et al*., 2022), CATH (Sillitoe *et al*., 2021), SCOP (Andreeva *et al*., 2020); **ligands**: ChEBI (Hastings *et al*., 2016), ChEMBL (Zdrazil *et al*., 2023), DrugBank (Knox *et al*., 2024); **publications**: ORCID (Haak *et al*., 2012), ISSN, journal names, PubMed, PubMed Central, DOI; **and standardised molecular weight** obtained by PDBe assemblies (Velankar *et al*., 2016).

### 2.2. Query system

The EMDB query system is powered by Apache Solr (Apache Software Foundation, 2025), utilizing the Lucene query language to enable a flexible and powerful filter mechanism for searching the database. This system allows users to combine multiple filters using boolean operators and range queries, providing high adaptability in constructing complex searches tailored to specific criteria. For example, users can filter entries by resolution ranges (e.g., resolution:[3 TO 5]) or apply time-based filters to select data deposited within a certain date range (e.g. deposition_date:[ 2024-01-01T00:00:00Z TO 2024-12-31T00:00:00Z]). Full documentation on how to use the EMDB search system and the list of available search fields can be found in the website (https://www.ebi.ac.uk/emdb/documentation/search, https://www.ebi.ac.uk/emdb/documentation/search/fields).

### 2.3. Implementation details

The implementation of the EMDB Chart Builder relies on a backend architecture composed of REST API endpoints developed using Django (Django Software Foundation, 2025) and Python. These backend services communicate directly with the Solr search engine to execute user queries, retrieve, and format the relevant metadata and validation information from EMDB and EMPIAR. On the front-end, Highcharts (Highsoft, 2025) is used as the visualization library to render interactive and customizable statistical plots. This architecture allows flexible data exploration with rapid responses, offering an intuitive user experience that does not require users to handle raw data or scripting directly.

The data underlying the Search System is updated on a weekly schedule, during the EMDB weekly release cycle that occurs every Wednesday at midnight UTC (wwPDB Consortium, 2024). For every new release and updated entries scheduled to the current weekly release, EMDB indexes the metadata and performs a full run of the Validation Analysis package, indexing the results in the Search System. The EMICSS pipeline is executed for every entry of the weekly archive release, with regular updates of the archive (Duraisamy *et al*., 2025), updating all the Solr search documents to the latest version of data from cross-referenced databases.

### 2.4. Reproducibility and usability

The EMDB Chart Builder promotes reproducibility and usability by providing open access through the EMDB website, ensuring that all users can freely explore, create, and share custom charts without restrictions, in line with EMDB’s open data policies. A key aspect of reproducibility is the ability to trace and adapt the visualizations that are already in use across the resource. To support this, most of the charts displayed on the EMDB statistics page and within search result statistics now include an “Edit” button that opens the corresponding plot directly in the Chart Builder. This feature allows users to examine the underlying configuration of predefined charts, use them as worked examples, and then adapt or extend them to address their own research questions. In addition, the tool guarantees consistency between chart visualizations and corresponding tabular output, enabling users to seamlessly switch between graphical and numerical representations for validation and deeper analysis. The Chart Builder is fully integrated into the existing EMDB website and can be accessed at https://www.ebi.ac.uk/emdb/statistics/builder.

## 3 Results

The Chart Builder provides an interactive user interface where users can generate custom visualizations from EMDB and EMPIAR metadata without the need for programming. The tool integrates a query-driven interface (Figure 1a) with multiple chart types, flexible axis definitions, and customizable data series, allowing users to explore archive statistics in a reproducible and transparent manner. In the following subsections, we demonstrate the main functionalities of this web based user driven visualisation and analysis tool.

**Figure 1.**
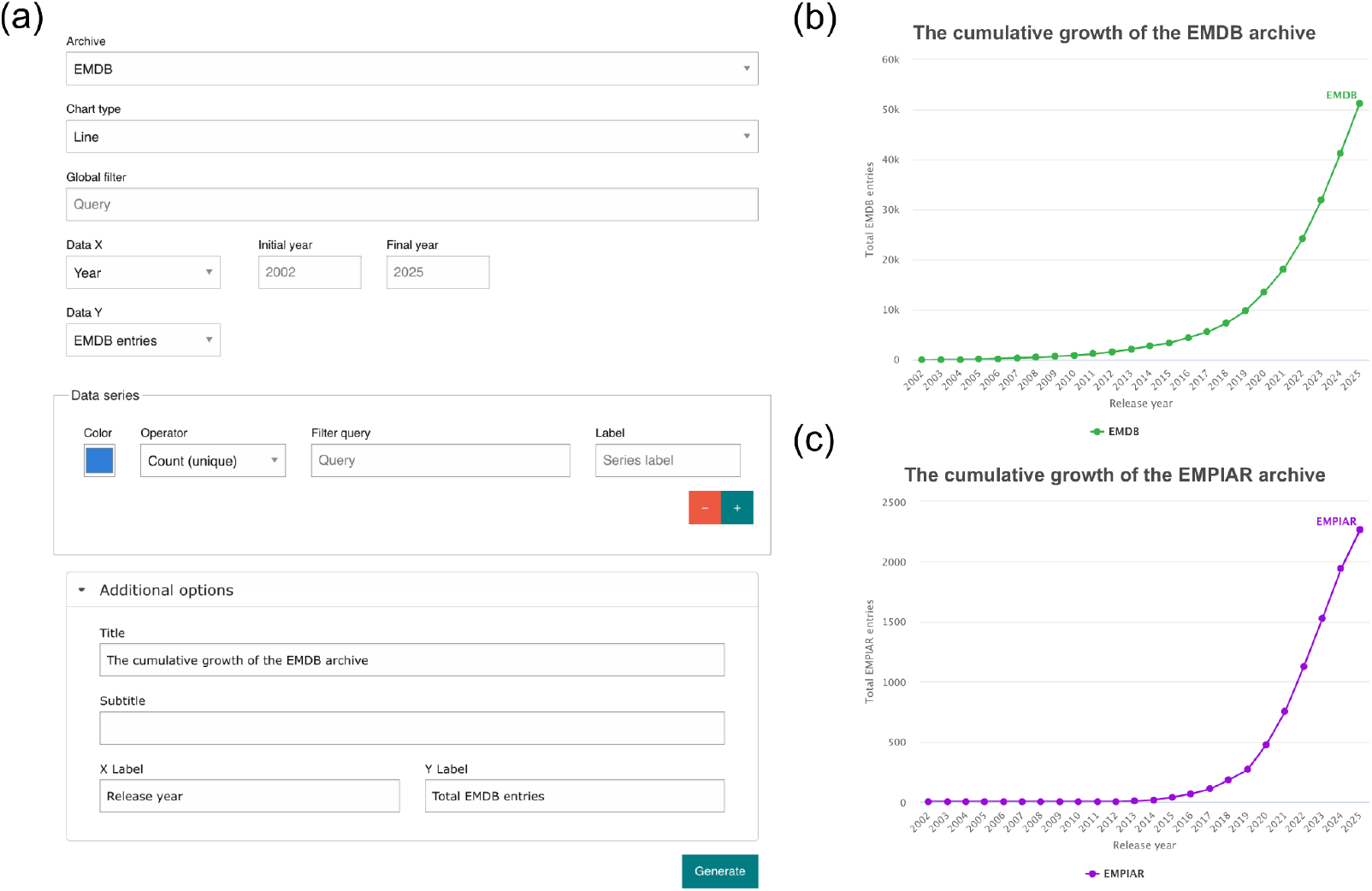
The basic (a) chart builder layout and two simple line plots generated from Chart Builder showing the cumulative growth of (b) the EMDB and (c) EMPIAR (b) archives.

### 3.1. Archive Selection

Users start by selecting the archive from which the data will be drawn: either EMDB or EMPIAR. The selection of the archive will change what search fields are available for use when building the chart. Figure 1 contains two graphs generated by the Chart Builder showing the growth of EMDB and EMPIAR archives on an annual basis.

### 3.2. Chart Types

Chart Builder supports a diverse range of visualization types to accommodate different data exploration and analytical needs. The tool includes standard formats such as line, area, bar and stream; statistical formats such as histograms, pie charts and scatter plots; as well as more specialized visualizations such as bubble plots, Venn diagrams and map charts.

#### 3.2.1. Bar and area charts

Bar and area charts can be displayed in stacked, unstacked, or percentage-stacked (Figure 2a) formats, making them suitable for analysing distributions and proportions across time or metadata categories.

**Figure 2.**
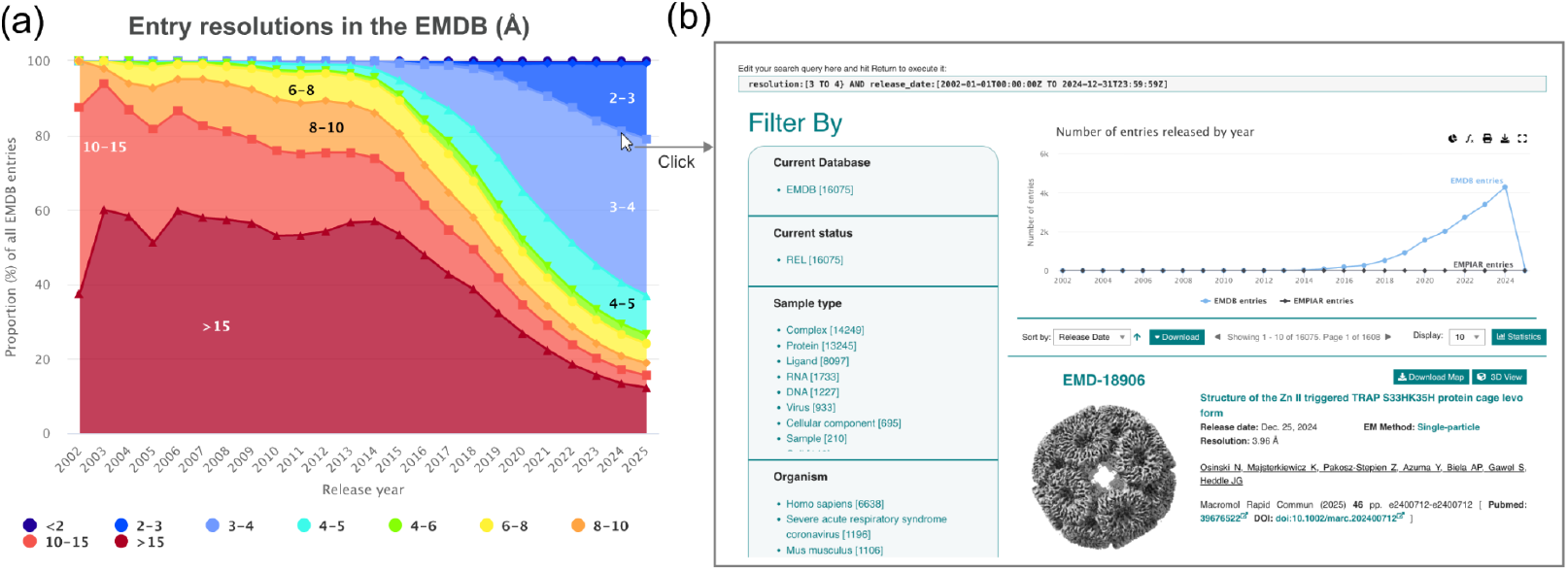
Area plots may be visualized as unstacked, stacked, or (a) stacked-percent, revealing resolution data distribution over time. User interaction with distribution based data series charts opens (b) search results pages representing these data.

Each chart is interactive and can be customized to display data series according to user-defined filters, colours, and statistical operators, such as maximum, minimum, average, count or unique. All charts can be interacted with, where hovering with the cursor will display an inspection of the data for that distribution data, and a click event will redirect to the EMDB search results (Figure 2b) for the corresponding data of that part of the chart. Most of the chart types share the same controls, some exceptions are listed below.

#### 3.2.2. Pie charts

Pie charts are limited to a single data series and provide a clear overview of proportional breakdowns (e.g., fraction of entries by EM method). The pie chart slices are defined by the dataset provided to the X-axis option, and the values are defined by the Y-axis option.

#### 3.2.3. Scatter plots

Scatter (2D and 3D) and bubble plots allow the exploration of relationships between numerical variables. Instead of defining x- and y-axes through categorical or temporal variables, users directly select numerical variables (search fields) from a dropdown (e.g., molecular weight, resolution, map size). Each point in these plots corresponds to an individual EMDB or EMPIAR entry and the click event will redirect to the respective entry page instead of the search results. Bubble and Scatter 3D plots further extend this by incorporating a third numerical dimension (z-axis), such as dataset size, resolution or other validation metrics, which determines the z-axis scaling (bubble size or point color) as seen in Figure 3.

**Figure 3.**
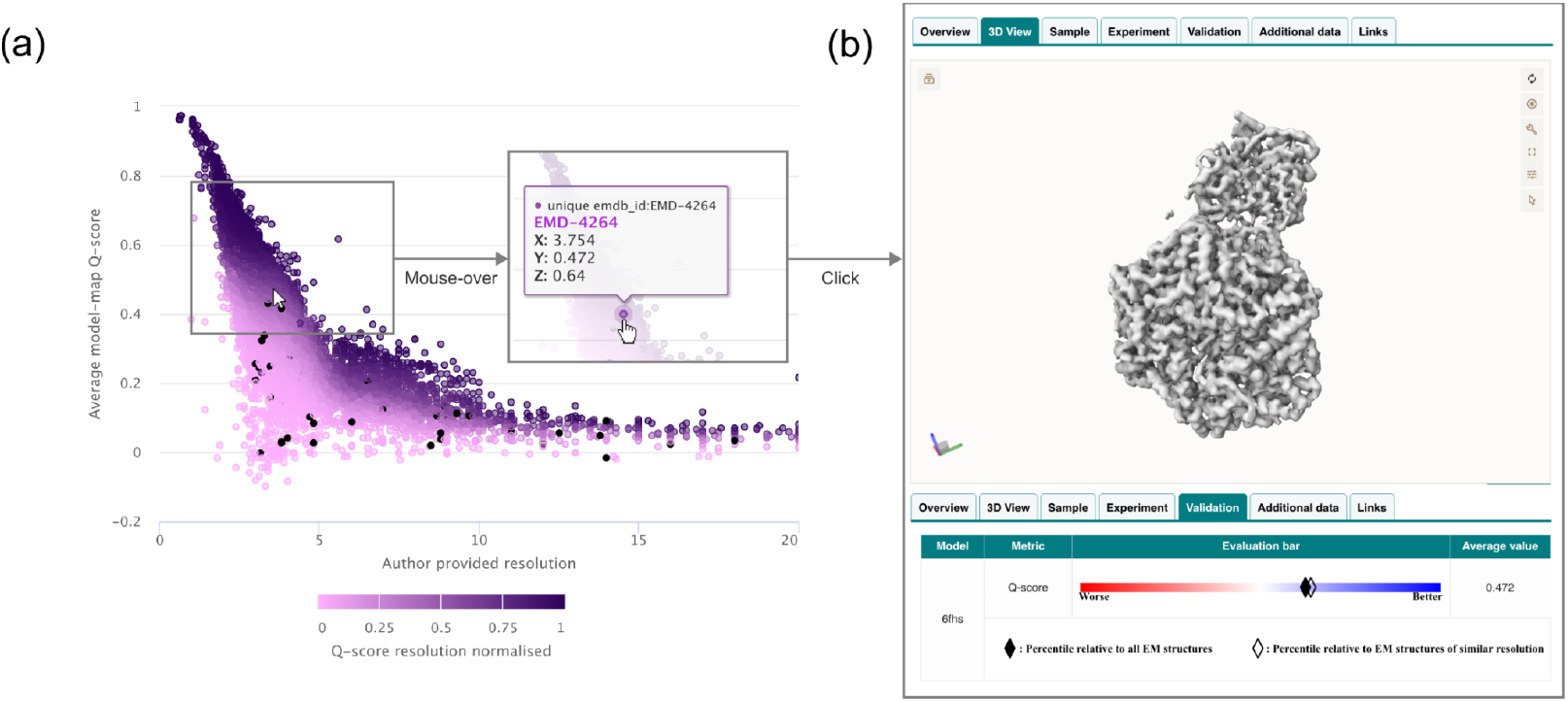
(A) Exploring a model-map fitness and resolvability measure (Q-score) and its resolution dependence for the entire EMDB archive. A third data type ranking Q-scores between 0-1 within resolution bins is used to color the data points. (B) Contributing data may be inspected for scatter plots where a click event on a data point will open the corresponding EMDB entry page.

**Figure 4.**
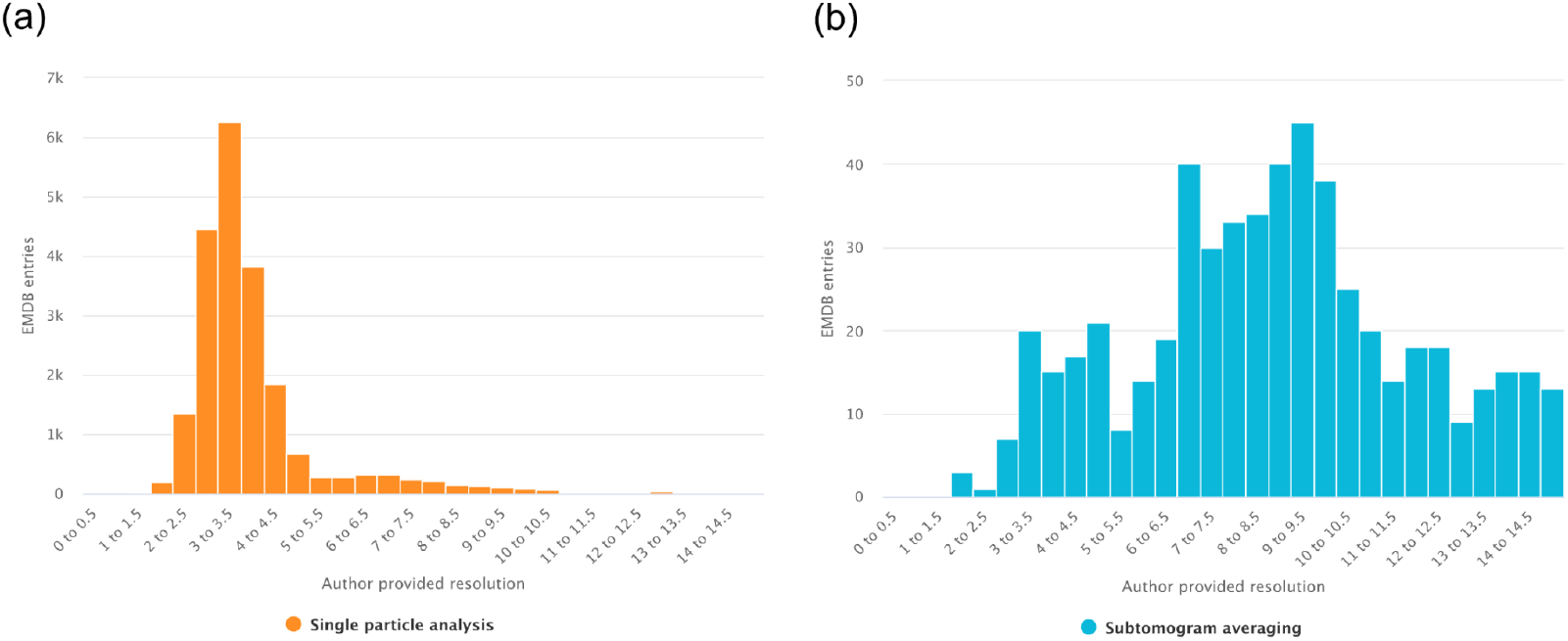
Evaluating the resolutions of 3DEM entries using global filters for structure determination method. (A) Map resolutions for entries determined by single particle analysis. (B) Map resolutions for entries determined by subtomogram averaging.

#### 3.2.4. Histograms

Histograms enable users to examine the distribution of numerical variables within EMDB or EMPIAR datasets. Similar to scatter plots, histograms require the user to select one numerical variable from the available search fields to define the X-axis (for example, resolution, map voxel size, or molecular weight). The frequency or count of entries within user defined value intervals is then represented along the Y-axis.

User-defined value intervals are defined for the histogram by specifying the start value, end value, and bin (gap) size, which allows control over the resolution and range of the displayed distribution. This flexibility makes histograms particularly useful for identifying trends, outliers, or biases in data distributions.

#### 3.2.5. Venn diagrams

Venn diagrams provide a unique way to visualize overlaps between sets defined by user-specified queries. In this plot, users also don’t define x- and y-axes but instead use the data series to define each set. Each data series corresponds to a set, and intersections illustrate entries that satisfy multiple conditions (e.g., entries in EMDB with fitted models in PDB, corresponding models in AlphaFold DB and raw data in EMPIAR). This feature enables users to investigate relationships between subsets of the archives in an intuitive manner.

#### 3.2.6. Geographic plots

Geolocation annotations are available for both EMDB and EMPIAR entries. For EMPIAR, we provide statistics for the principal investigator’s institutional affiliation’s country and continent, since its information is available in the EMPIAR metadata. For EMDB, we provide access to wwPDB sites (deposition and annotation) and grant country, both collected from the deposition; and we also have four enriched fields collected from the Europe PMC: corresponding author country, continent, city and institution (Rosonovski *et al*., 2024). These will only include EMDB entries with an associated publication that is in open access format. Depending on the type of data, the geographic plots can be coloured as heatmap or bubbles. In the figure 5 we show both examples, the number of EMDB entries released per continent as a heatmap, and the number of EMDB entries released by institutions on the Research Organization Registry (Research Organization Registry, 2024) visualized as a bubble map.

**Figure 5.**
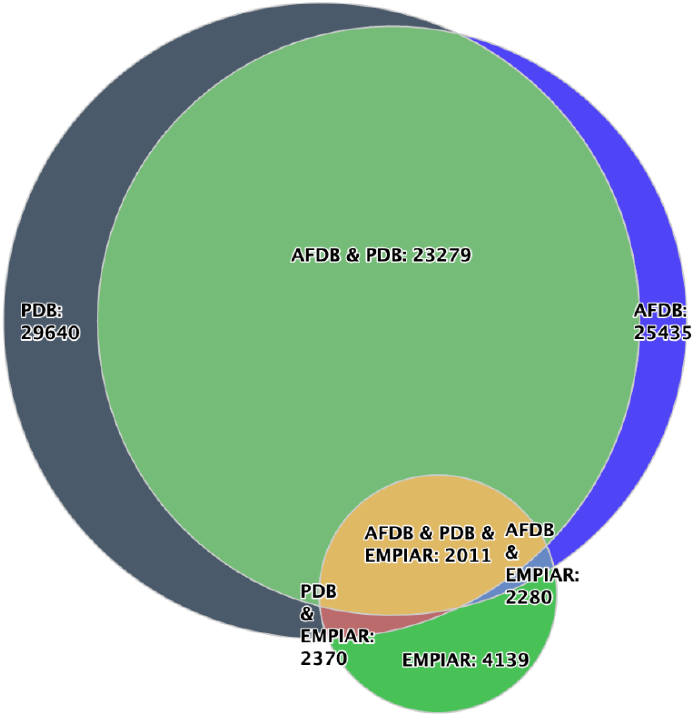
A venn diagram representing EMDB entries where data is available in complementary databases such as EMPIAR, PDB and AlphaFold Database.

### 3.3 Global filter

Before generating a chart, users can apply a global filter to restrict the entire dataset that will be provided to the chart. This filter is expressed as a query in the EMDB query language i.e. ***structure_determination_method:”singleParticle”***. A global filter can target any available metadata field, such as reconstruction method, release year, or sample type. By narrowing the dataset at the outset, researchers can focus their analysis on a specific subset of interest. For example, a query restricted to single-particle reconstructions allows users to investigate resolution trends only within that method, while excluding tomography or subtomogram averaging entries.

### 3.5. X-Axis Options

The independent variables (x-axis) of a chart can be defined in three different ways, enabling both temporal and categorical analyses:

Yearly - Users may specify a start and end year to visualize trends over time. This is particularly effective for examining archive growth, such as the steady increase in high-resolution structures deposited since 2015 as seen in Figure 2.

By Experimental Metadata – A categorical variable (e.g., EM method, sample type, or microscope model) can be chosen as the x-axis, with all categories automatically included in the visualization.

This option enables comparisons across experimental conditions, for example showing the relative contribution of different microscopes to deposited structures.

By Custom Queries – Users can define a set of queries manually, with each query corresponding to one label on the x-axis. This provides maximum flexibility, allowing the creation of user-defined categories such as a distinct set of software as seen in Figure 6.

**Figure 6.**
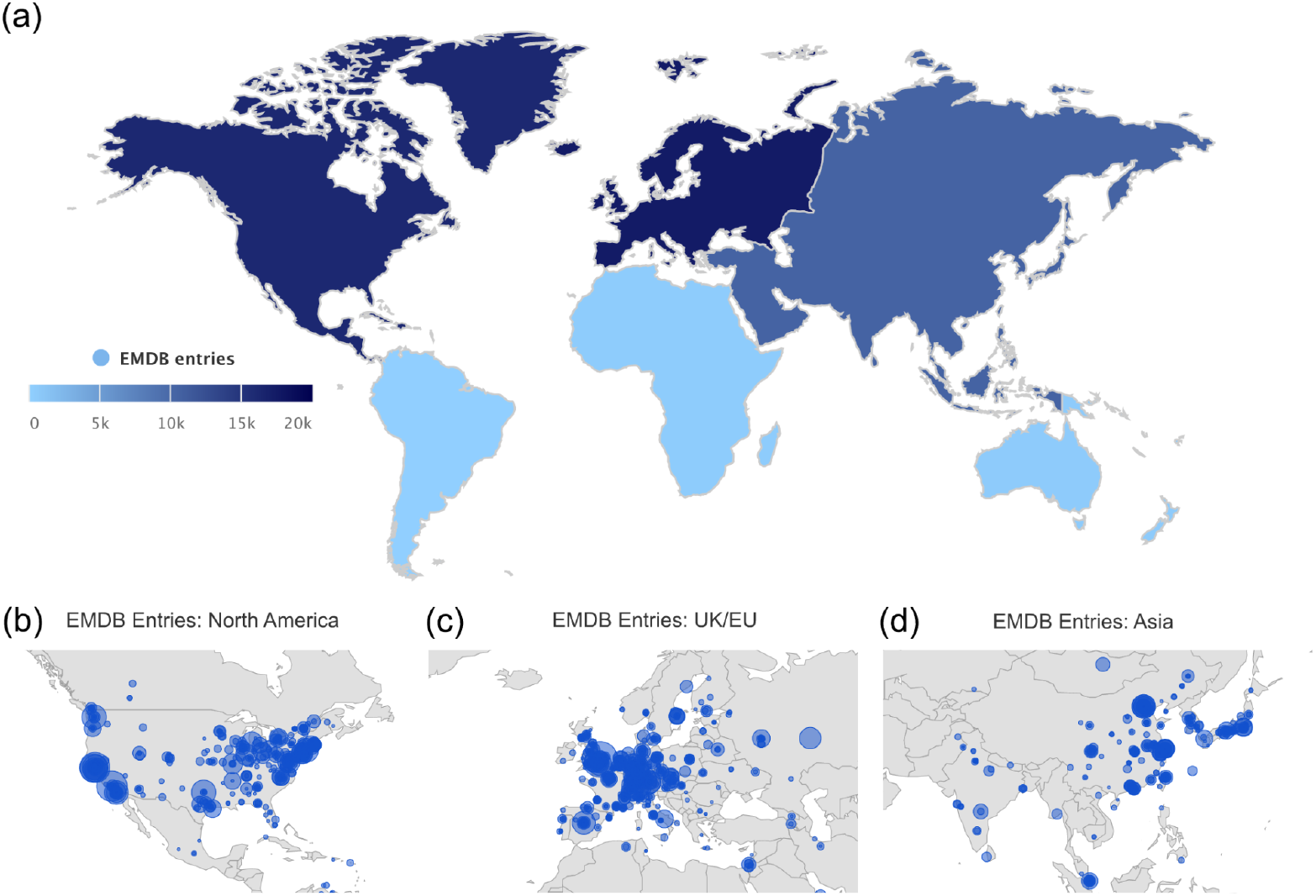
Two geographic plots showing the number of EMDB entries released. (A) per corresponding author country (B) per corresponding author institution for the major three data producing continents.

Together, these options allow users to construct temporal, categorical, or bespoke groupings, making it possible to address a wide range of exploratory questions with a single interface.

### 3.6. Y-Axis Options

By default, the dependent variables (y-axis) are displayed as the number of EMDB/EMPIAR entries. However, users can select alternative variables, such as for EMDB: number of authors, number of fitted models, number of associated raw data entries in EMPIAR, number of masks, number of additional maps, resolution, number of publications, molecular weight, atom inclusion, Q-score, map size, and frames per image; and for EMPIAR: number of authors, number of associated EMDB entries, number of Publications, dataset size, and number of images. Figure 6 shows two examples using distinct data Y options: count of publications and minimum/average/maximum molecular weight.

### 3.7. Data Series

The Chart Builder allows the inclusion of multiple data series within a single plot, enabling comparative and multi-dimensional analyses. Each data series can be customized with a user-defined color, statistical operator (count, sum, average, maximum, or minimum), and an optional filter query to refine the subset of data being represented. Users may also assign custom labels, which are displayed in the chart legend to ensure clarity in interpretation. An example on how data series can be used in this application is shown in Figure 7.

**Figure 7.**
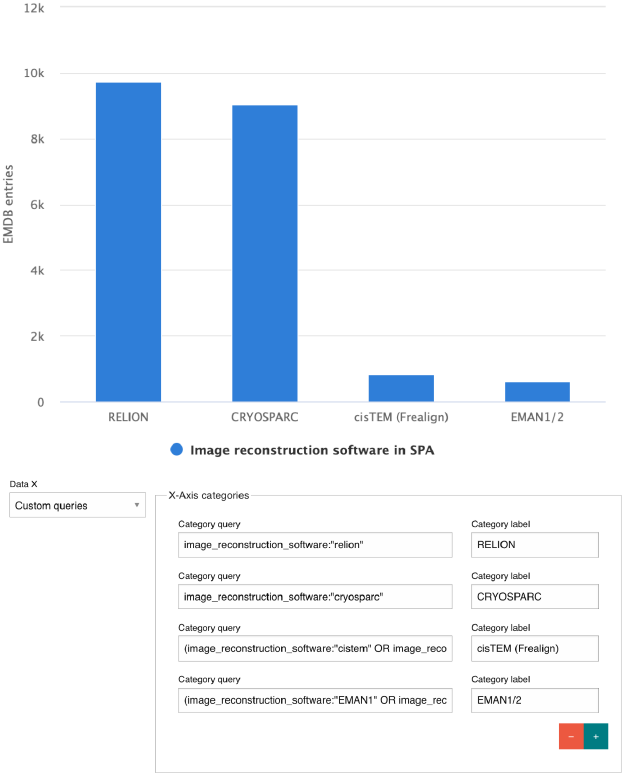
Using custom queries to compare the image reconstruction software usage in EMDB entries from Single Particle Analysis.

**Figure 8.**
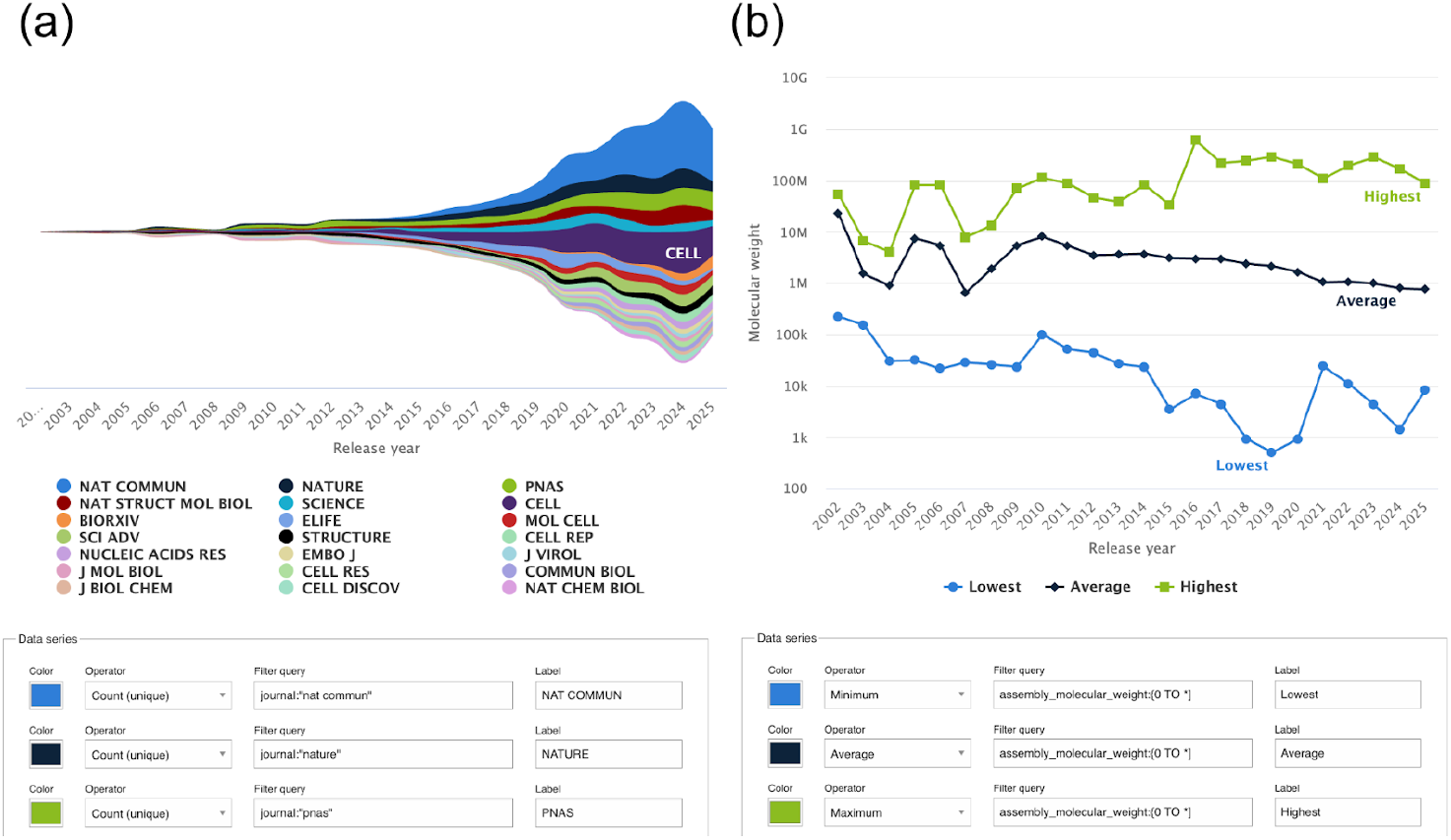
Two graphs using multiple data series and non-standard data Y options. The first shows journal trends in EMDB over the years as a Stream plot, with multiple data series composed by small queries filtering each data series to specific journals. The second shows molecular weight trends in EMDB, using three data series without filtering, but instead playing with the operators to show maximum, average and minimum value per year.

### 3.8. Chart Customisation and Export

The Chart Builder provides a range of interactive and customization options that enhance both the usability and adaptability of the visualizations. Once generated, charts remain fully interactive: users can dynamically switch between display modes such as stacked, unstacked, or percentage views (for bar and area plots – Figure 2) and toggle between linear and logarithmic scales (Figure 6) to effectively represent data spanning multiple orders of magnitude. Data series can also be interactively managed; clicking on a legend item allows users to toggle individual series on or off, facilitating clearer comparisons between subsets of data. These features are accessible via a menu at the top right of charts, the buttons of which are chart specific and summarised in table 1.

**Table 1.**
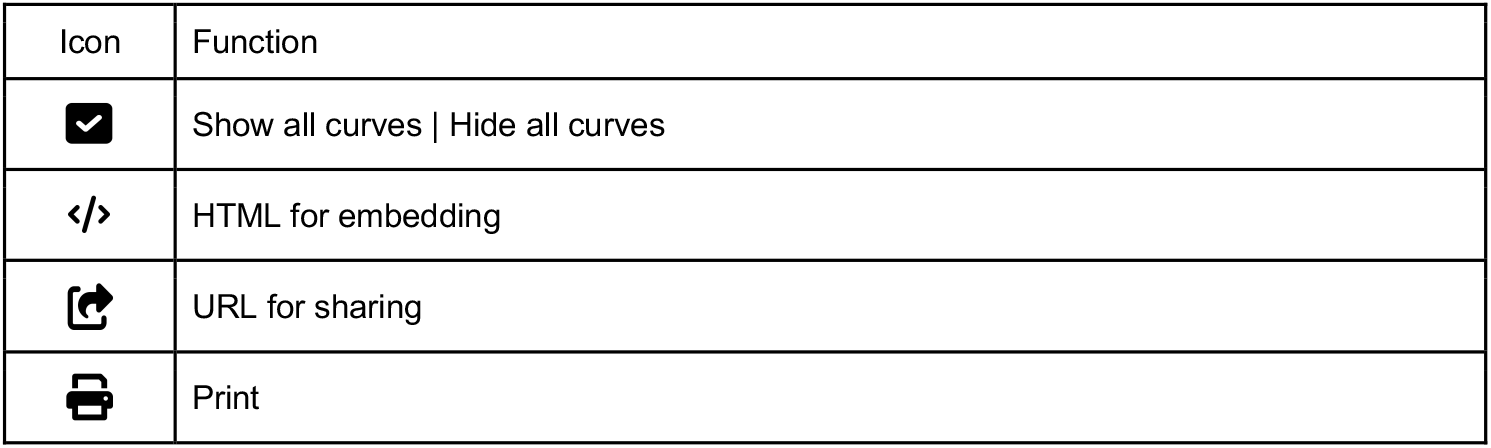
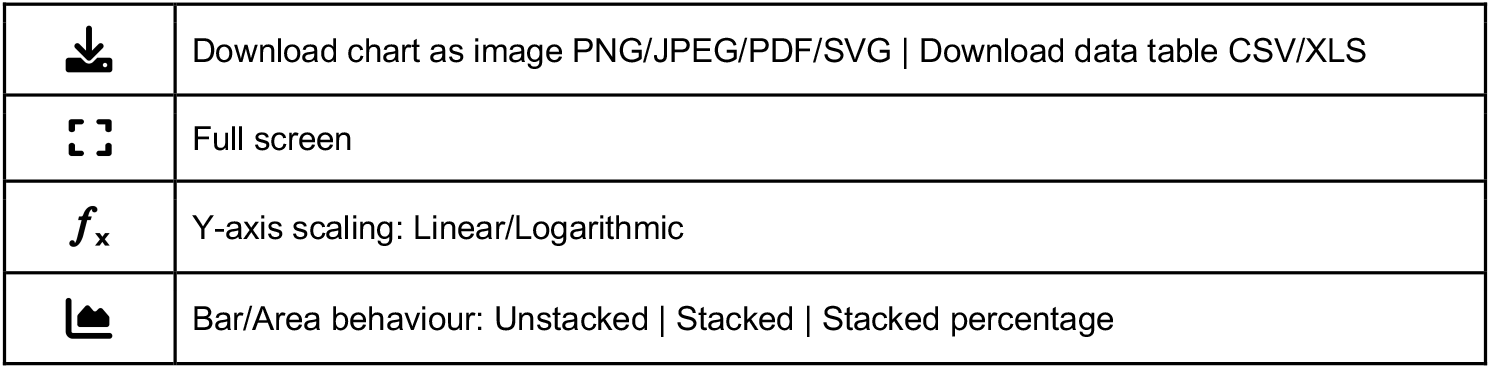
Descriptions of the data export and chart modification.

Additional interactive features support exploration and validation of the dataset which has been developed for plotting in a chart. Hovering over chart elements reveals precise numerical values relevant to that data point or series. While clicking on a data point in a scatter plot opens that specific entry for exploration. For the chart types histogram, pie, bar, when a chart section is clicked it opens a new window with search filters enabled to show results from that section of the chart. Export functionality is versatile by allowing the generation of shareable URL links to the customized chart builder page itself, generating HTML code for chart embedding, printable charts, and downloadable figures in multiple formats, including high-quality images (PNG, JPEG, PDF or SVG). Full screen views are suitable for presentations with support for granular customization for presentation and reporting including; Titles, subtitles, and axis labels directly editable within the interface. Finally, for further external data analyses, tabular files (CSV or XLS) of the data generating the current plot may be downloaded.

## 4 Discussion

The EMDB Chart Builder was developed to enable the statistical exploration of the expanding 3DEM landscape through intuitive, interactive configurable visualisations rather than scripts or specialist tools. As EMDB and EMPIAR continue to grow in size and complexity, tools are required to understand the archive at a glance to assess emerging patterns or contextualise deposited entries within broader community trends. The Chart Builder addresses this by leveraging the EMDB Solr search system into a visual, user-driven discovery environment where a broad user base can build, adapt, and share statistical summaries of the 3DEM structural biology archives. As a major step forward from the original fixed EMstats plots, the Chart Builder allows users to test unique hypotheses through configurable filters and metadata combinations, or refining existing EMDB curated visualisations. Lastly, these capabilities directly align with FAIR principles: they make data easier to find, easier to access, interoperable across metadata sources, and readily reusable through persistent URLs and export options.

A central strength of the tool lies in the harmonisation of metadata from multiple sources—including depositor-supplied fields, EMICSS cross-references, and Validation Analysis metrics—into a single visual interface. This integration empowers the community to develop rich data driven insights: how 3DEM data properties inferred by validation are distributed, which software pipelines are shaping current practice, how molecular weights or resolutions vary across methods and time, or how data production is geographically distributed. User-level interactions support exploring these insights; hovering exposes metadata and clicking linking directly back to underlying entries or archive searches.

Figure 9 presents a use case where a data visualization may be used to identify a cluster of the EMDB archive with properties valuable to molecular modelling methods development. Further examples in this manuscript illustrate how visual analytics can support monitoring archive growth, exploring model–map fit, examining software usage and molecular weight trends, analysing cross-archive relationships, and mapping global 3DEM methodology trends. Collectively, these use cases demonstrate how visual tools can reveal clusters, anomalies, or methodological signals that are useful for benchmarking, training-set creation, and methodological innovation.

**Figure 9.**
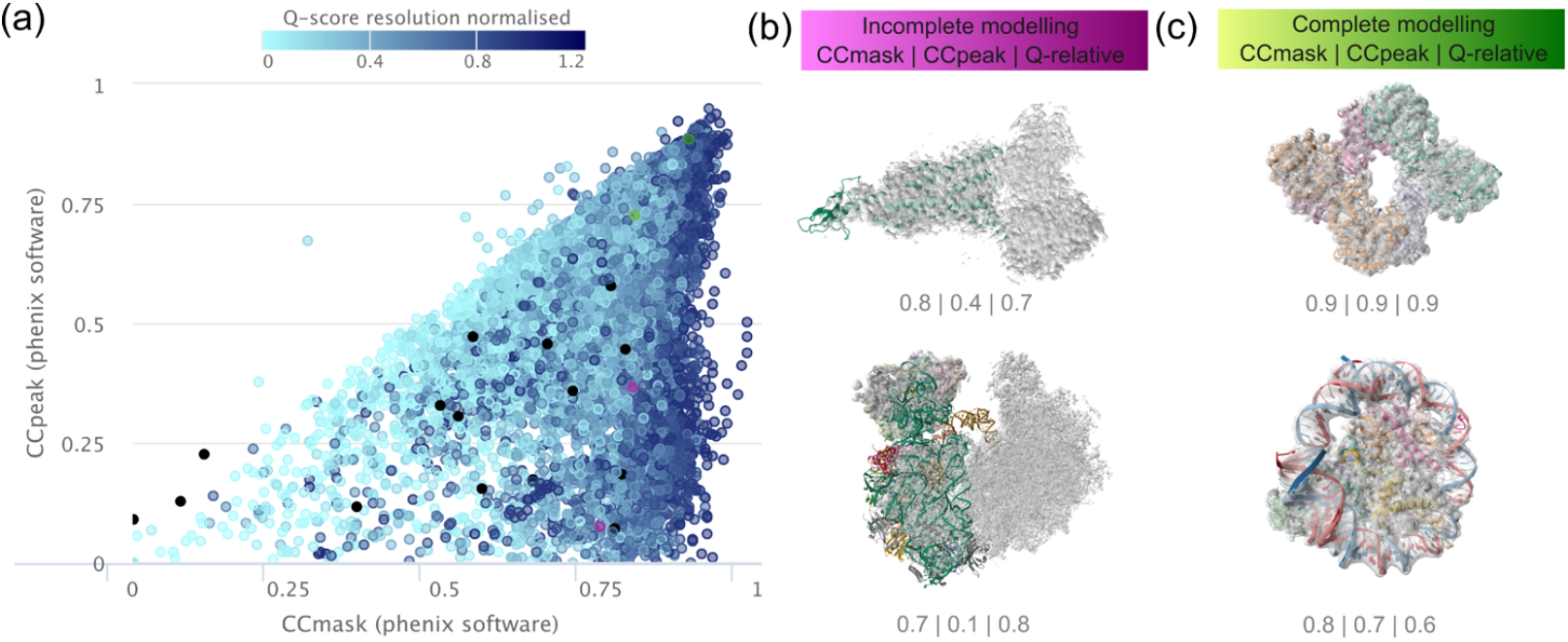
A use-case demonstrating a dataset discovery approach with the use of chart builder visualisations. Entries are identified (A) in a cluster in the lower-right quadrant with high CCmask/low CCpeak which may represent entries with unmodelled 3DEM density (Afonine et al., 2018). (B) Entries with opportunity for further modelling (EMD-23275/PDB 8G4I and EMD-29721/PDB 8G4I) from this cluster are shown to illustrate the potential of this use case. In comparison (C) entries with complete modelling are found where CCmask **≈** CCpeak (EMD-15028/PDB 7ZYY and EMD-X/PDB X).

Altogether, the Chart Builder aims to support users by lowering barriers to data exploration and enabling researchers to interact with the archive in ways that are visual, open, and collaborative. It provides a modern platform for transparent and reproducible exploration of EMDB and EMPIAR, helping make electron microscopy data more accessible and interpretable, and supporting the need for data-driven insight, innovation, and discovery.

## 6 Author Contributions

Neli Fonseca (Conceptualization, Formal Analysis, Software, Writing-original draft, Writing–review & editing). Gerard J. Kleywegt (Conceptualization, Supervision), Ardan Patwardhan (Funding acquisition), Matthew Hartley (Supervision, Writing–review & editing) and Kyle L. Morris (Conceptualization, Supervision, Writing–review & editing, Funding acquisition). All authors have read and agreed to the published version of the manuscript.

## 8 Acknowledgements

We thank Matt Jeffryes and Melissa Harrison in EuropePMC (EPMC @ EMBL-EBI) for discussion around EPMC API endpoints to provide geographical annotations to EMDB entries. We thank the wwPDB in the worldwide collaborative effort to archive structural biology data and the OneDep team for continued team work to deliver this vision. We acknowledge the use of ChatGPT (OpenAI) as a tool to assist with spell checking, grammatical corrections and editorial assistance during the preparation of this manuscript. All intellectual content, methods, codes, interpretations, and conclusions are the sole responsibility of the authors.

## 9 Data Availability Statement

The datasets analyzed for this study can be found at www.emdatabank.org. All chart visualizations are viewable via persistent URL links found in the supplemental materials. All chart images used in this work are available for download from the persistent chart URLs. Individual EMDB entries that have been used for illustrative purposes are available in the EMDB under the accession codes; EMD-4264, EMD-15028, EMD-22692, EMD-23275 and EMD-29721. All entry images used in this work are available for viewing and download from www.emdatabank.org entry page 3D viewer tabs, using the web-based Mol* plugin (Sehnal et al., 2021).

---

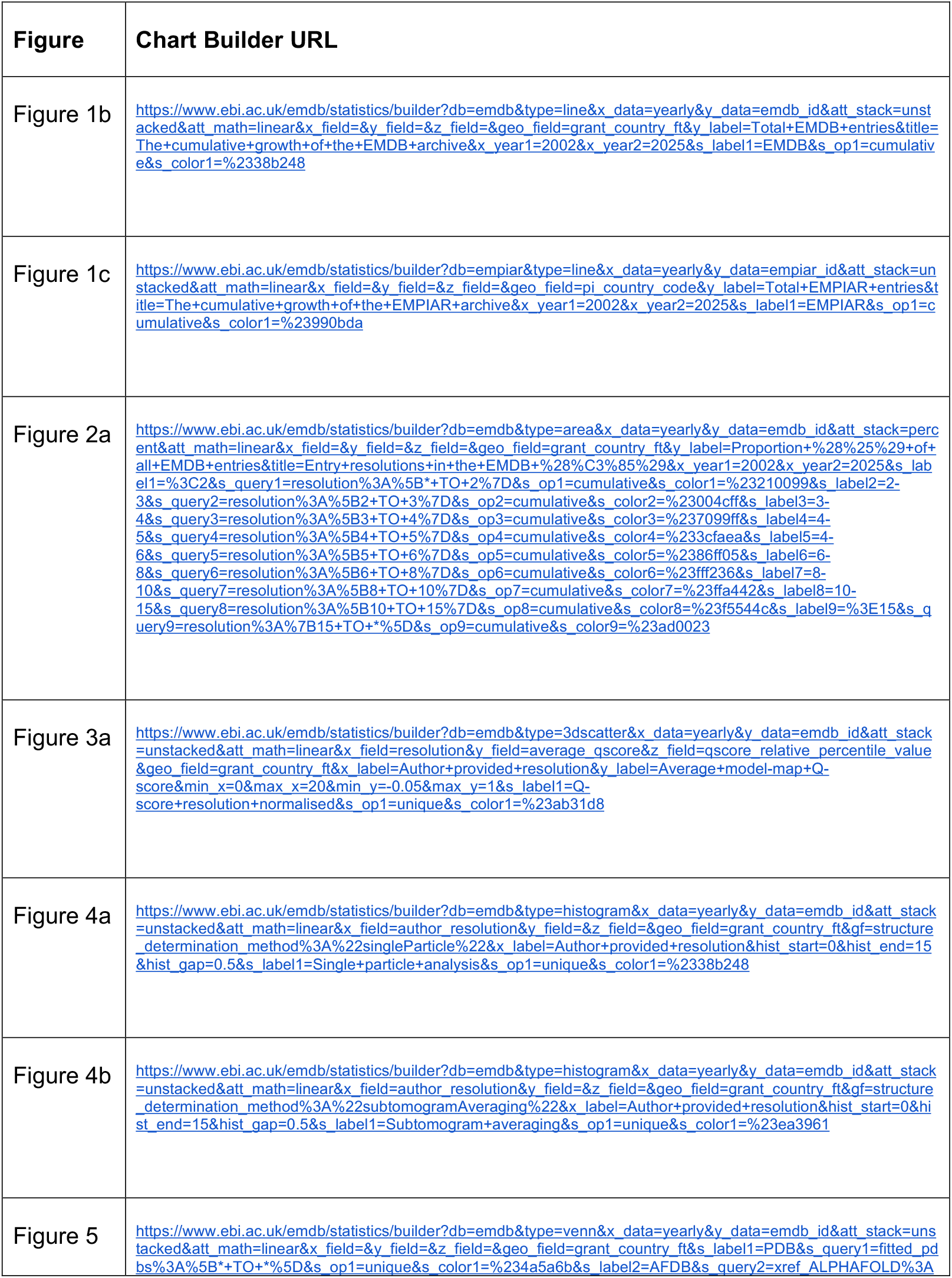
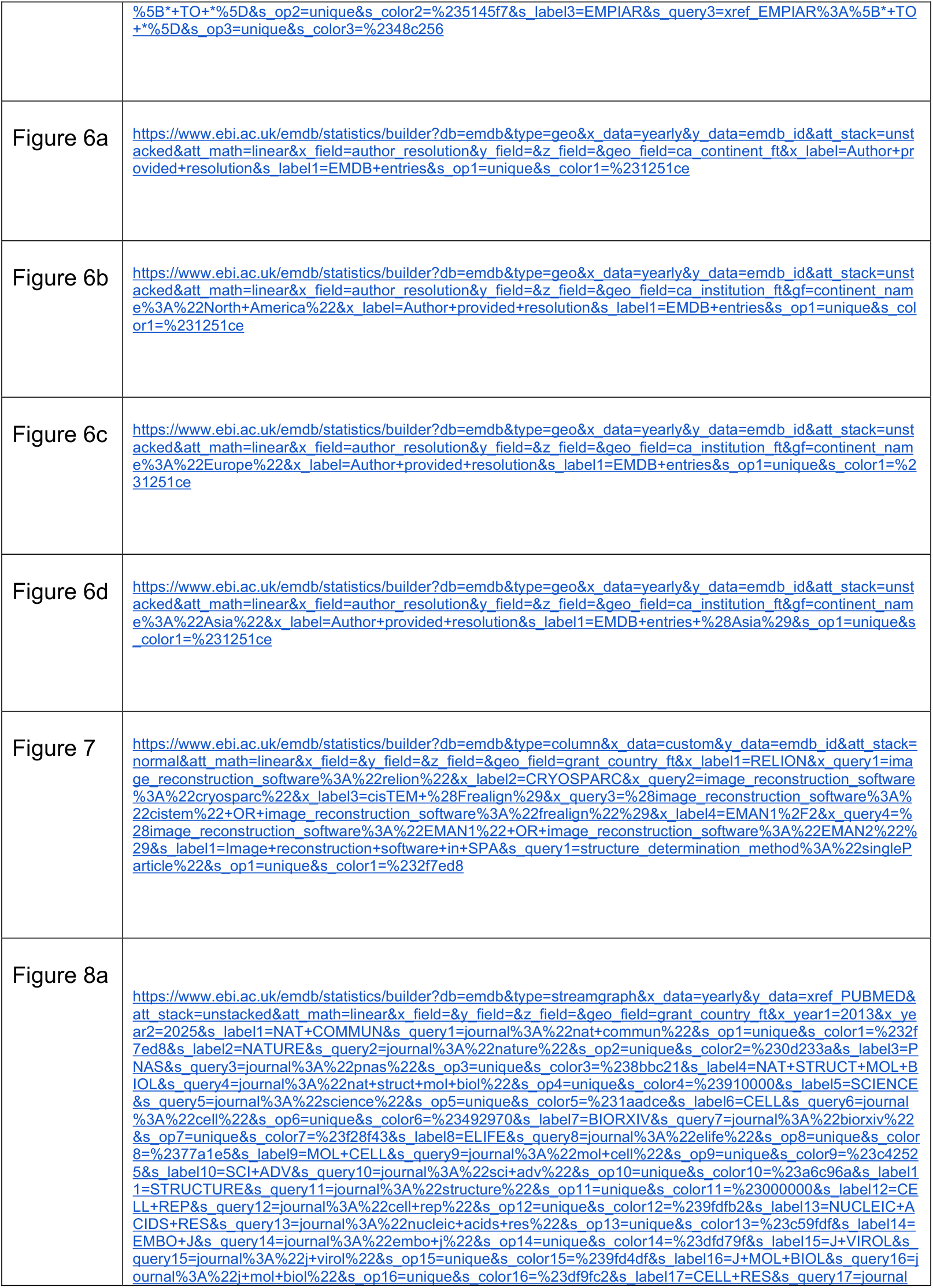
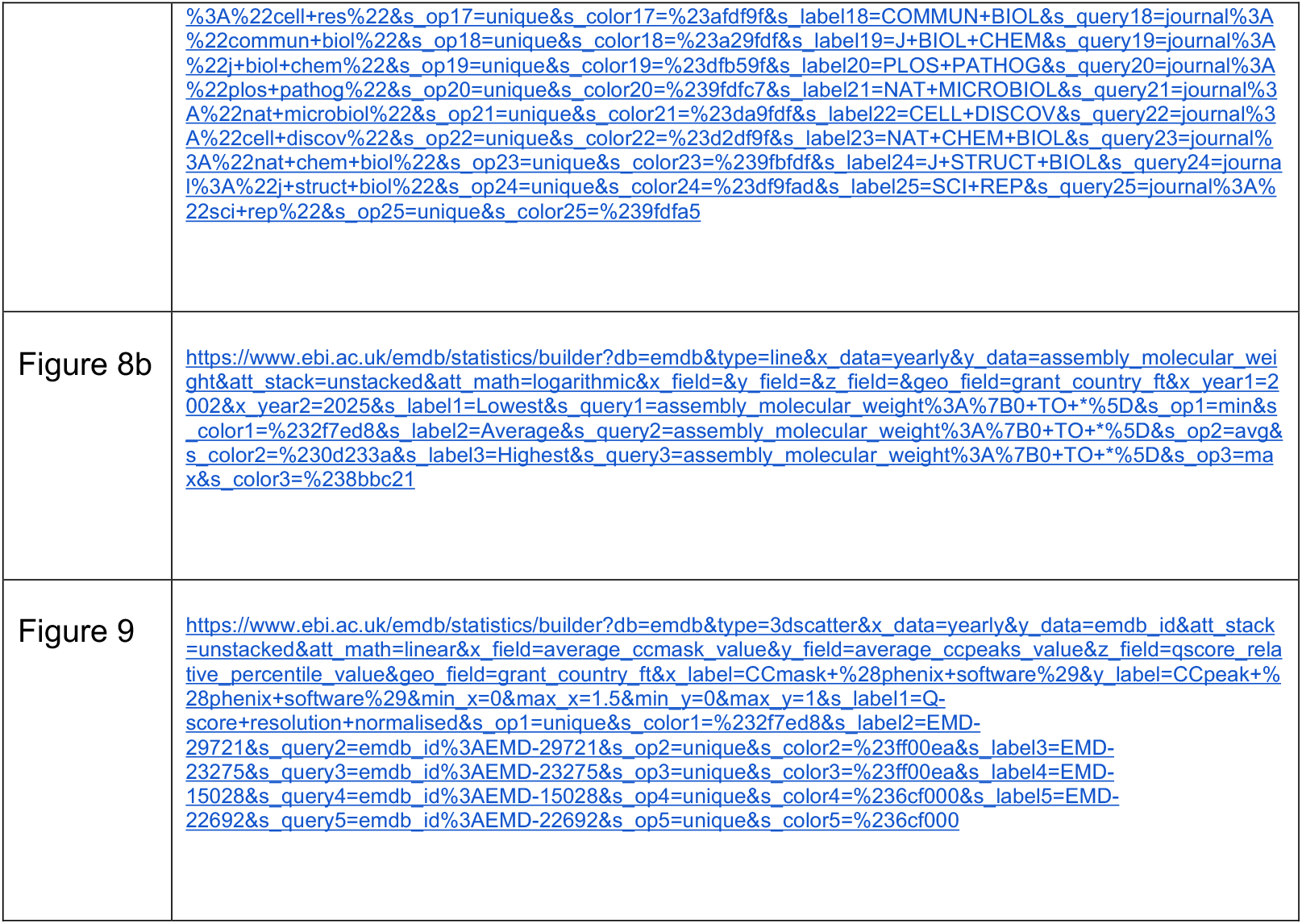

